# Bioactivity, chemical profiling, and 16S rRNA-based phylogeny of haloalkaliphilic *Nocardiopsis sp. GhM-HA-6* isolated from the Gulf of Khambhat

**DOI:** 10.1101/2021.07.05.451136

**Authors:** Nisha D. Trivedi, Jignasha T. Thumar

**Affiliations:** Department of Microbiology, School of Science, RK University, Rajkot 360020, Gujarat, India; Department of Microbiology, Government Science College, Gandhinagar-382015, Gujarat, India

**Keywords:** marine actinomycetes, haloalkaliphilic, antimicrobial activity, gas chromatography-mass spectroscopy, 2, 4-bis (1, 1-dimethylethyl)-Phenol

## Abstract

**Objectives:** Actinomycetes are well known sources of antibiotics, however; recently the focus of antimicrobial research has been turning towards actinomycetes of extreme environments. Therefore, present work would highlight the isolation, identification and characterization of antimicrobial metabolites produced by marine haloalkaliphilic actinomycetes.

**Methods:** Saline soil sample was collected from Ghogha coast (Gulf of Khambhat), Bhavnagar, Western India. Isolation was carried out using selective media while identification was done based on morphological, cultural and molecular characterization. The antimicrobial potential was checked by spot inoculation method. Optimization was carried out by the one variable at a time (OVAT) method. The antimicrobial compounds were extracted using ethyl acetate and characterized by GC-MS.

**Results:** The haloalkaliphilic actinomycetes *Nocardiopsis sp.* GhM-HA-6 was isolated from saline soil of Ghogha coast using starch agar with 10% w/v NaCl and pH 9 and was identified as *Nocardiopsis sp.* based on morphology, cultural characteristics and 16S rRNA phylogenetic analysis (NCBI Genbank Accession number: KF384492). The organism showed antimicrobial activity against five Gram positive and three Gram negative bacteria while the isolate didn’t show any antifungal activity. Results of optimization showed that the highest production of antimicrobial compounds was obtained using starch broth with 0.5% w/v starch, 1% w/v yeast extract, 10% w/v NaCl and pH 9. GC-MS analysis of ethyl acetate extract of the isolate showed presence of total 18 compounds including various antimicrobial compounds like 2, 4-bis (1, 1-dimethylethyl)-Phenol, various types of alkanes and their derivatives.

**Conclusion:** Haloalkaliphilic actinomycete *Nocardiopsis sp.* GhM-HA-6, from a rarely explored marine habitat, can be a source of antimicrobial compounds with the novel biotechnological applications.

## 1. Introduction

Actinobacteria are well known for their ability to produce valuable secondary metabolites such as antibiotics and antimicrobial substances. Amongst genera of actinobacteria, *Nocardiopsis* genus was primarily described by Meyer in 1976 and was placed in the class of Actinobacteria; subclass Actinobacteridae; order Actinomycetales and family Nocardiopsaceae. Rainey *et. al*., (1996) defined a new family referred as *Nocardiopsaceae* considering phylogenetic position, morphological features, and chemotaxonomic properties of *Nocardiopsis sp.* This genus includes Gram positive, aerobes, non-acid fast with catalase positive properties (Cook and Meyers, 2003). Studies by Kroppenstedt and Evtushenko (2006) showed various features of genus *Nocardiopsis* which includes the presence of meso-2, 6-diaminopimelic acid, but lack of diagnostically important carbohydrates in cell wall structure, absence of madurose or nocardomycolic acids in whole cell hydrolysates and high GC content in their genomes. Actinobacteria are widely found in soil and physiologically extreme environments such as desert, saline, hypersaline, and alkaline origins. Particularly, *Nocardiopsis* species are often found from hypersaline environments such as soda lakes and salterns along with *Streptomyces* species (Mwirichia et. al., 2010; Jose and Jebakumar, 2012). It is believed that *Nocardiopsis* genus might be playing a mediating role in breakdown of naturally occurring complex polymers, these species mainly produce extremozymes, compatible solutes, and surfactants (Bennur et. al., 2015). Genus *Nocardiopsis* has been reported to possess gene clusters for polyketide synthases (PKSs) and non-ribosomal peptide synthetases (NRPSs) and additionally many of the natural isolates have also demonstrated capability antibiotic production (Zitouni et. al., 2005; Meklat et. al., 2011; Becerril-Espinosa et. al., 2013; Romanenko et. al., 2013). *Nocardiopsis sp.,* often secretes important antimicrobials including polyketides, phenzines, quinoline alkaloids, terphenyls, proteins, thiopeptides, and amines (Bennur et. al., 2015). Marine isolates of *Nocardiopsis sp.* are often reported to produce cyclic hexapeptides which render antimicrobial activity (Wu et. al., 2013). The production of a novel antibiotic 3-trehalosamine was reported from soil-derived *N. trehalosei* by Dolak et. al., (1980). The study also reported production of dopsisamine and thiopeptide antibiotics by *N. mutabilis* and *Nocardiopsis sp.* TFS65-07, respectively (Takahashi et. al., 1986; Engelhardt et. al., 2010a).

Among the nine maritime states in the Indian peninsula, only a few states have been extensively covered for the study of marine actinobacteria for antagonistic properties against different pathogens (Sivakumar et. al., 2007). The Gulf of Khambhat is a coastal region situated in the Bhavnagar District of the Indian State Gujarat, which covers approximately 3120 km^2^ area of mudflat with some rocky (sandstones) intertidal area and a volume of 62,400 million m^3^ (Ramnathan et. al., 2002). Alang and Ghogha coasts of Gulf of Khambhat, India are relatively less explored with respect to antagonistic properties of actinomycetes. Ghogha is a small town situated at the mid-western shore of the Gulf of Khambhat, covering about 4 km long area (21°40’32” to 21°41’18” N and 72°17’5” to 72°16’48 E) in Bhavnagar, Gujarat with unique characteristics of having sandy supratidal zone, rocky-muddy middle intertidal zone and highly muddy lower intertidal zone (Solanki et. al., 2016). The uniqueness of this region is its salinity and alkalinity, which harbors various unidentified, unique haloalkaliphilic bacterial species that can potentially produce secondary metabolites. This highlights the urgent need for exploration of untapped location for novel antimicrobial antagonist producers. In the light of this knowledge, the present study was intended to isolate, screen and characterize the antimicrobial metabolite producing haloalkaliphilic actinomycetes from the Gulf of Khambhat.

## 2. Materials and methods

### 2.1 Isolation

The haloalkaliphilic actinomycete, *Nocardiopsis sp.GhM-HA-6* was isolated from saline soil of Ghogha coasts (Gulf of Khambhat), Bhavnagar (Longitude: 72° 11’ E and latitude: 21° 46’ N), India and treated with calcium carbonate and Ringer’s solution. Approximately 10-15g of wet soil sample was mixed with a pinch of calcium carbonate and was dried at 75°C for overnight. Next day, 1g of soil sample was added to 10ml of Ringer’s solution. From the supernatant, 0.1 ml of the sample was spread separately on starch agar (Hi-Media, India), containing 10% w/v NaCl and pH 9. The plates were incubated at 30°C; on sixth day a typical chalky white colony was appeared, the colony characteristics were noted and Gram’s staining was performed. On seventh day, colony was picked up and re-streaked on starch agar slants supplemented with 10% w/v NaCl and pH 9 to ensure the purity of the colony (Chakrabarti, 1998). The culture was maintained at 4°C.

### 2.2 Molecular Identification of Isolate by 16S rRNA

The molecular identification of the isolate GhM-HA-6 was carried out by 16S rRNA gene sequencing. The 16S rRNA gene was amplified using universal primers 518F 5’ccagcagccgcggtaatacg3’ and 800R 5’taccagggtatctaatc3’. PCR products were purified and sequenced. The resultant sequences aligned within the NCBI database (National Centre for Biotechnology Information) using BLASTN. The phylogenetic tree was constructed using neighbor-joining with Kimura 2-state parameter and pairwise-deletion model analysis implemented in the program MEGA software version X (Kumar et. al., 2018) and also evaluated further in a bootstrap analysis of 1,000 replicates. The 16S rRNA gene sequence of the isolate has been submitted to NCBI, GenBank, Maryland, USA.

### 2.3 The Antimicrobial Potential

Antimicrobial activity of *Nocardiopsis sp.GhM-HA-6* was checked by spot inoculation method (Kumar et. al., 2010) using starch agar (10% w/v NaCl, pH 9). The spore suspension of the isolate was spotted on the medium and incubated at 28°C until sporulation. The test organisms, procured from MTCC, Chandigarh (ATCC equivalent), were used to check antimicrobial activity. These included Gram-positive organisms: *Bacillus subtilis* (MTCC 441)*, Staphylococcus aureus* (MTCC 737)*, Micrococcus luteus* (MTCC 106)*, Staphylococcus epidermidis* (MTCC 3615), *Bacillus megaterium* (MTCC 428)*, Bacillus cereus* (MTCC 430); Gram-negative organisms: *Pseudomonas aeruginosa* (MTCC 424), *Escherichia coli* (MTCC 443), *Enterobacter aerogens* (MTCC 8560), *Serratia marcescens* (MTCC 97), *Shigella flexneri* (MTCC 1457), *Salmonella enteric typhimurium* (MTCC 98), *Proteus vulgaris* (MTCC 1771), *Klebsiella pneumonia* (MTCC 3384) and three fungi: *Aspergillus niger* (MTCC 282), *Fusarium oxysporum* (MTCC 284), *Candida albicans* (MTCC 227). The test organisms were grown in nutrient broth at 37°C for 24 hours. The molten nutrient agar, with 1% activated test culture, was poured on the sporulated *Nocardiopsis sp.GhM-HA-6.* After the incubation of 24 hours at 37°C, the zone of inhibition was measured for each test organism.

### 2.4 Optimization of Growth Conditions for Antimicrobial Compound Production

To achieve the highest production of antimicrobial compounds, effect of various growth conditions were studied by the one variable at a time (OVAT) method. The effect of various media (starch broth, starch casein broth, actinomycetes broth, glucose aspargine broth, glucose glycine broth, soybean malt broth and complete broth with 10% w/v NaCl and pH 9), effect of starch as a carbon source (0-1.5% w/v), effect of yeast extract as a nitrogenous source (0-1.5% w/v), effect of NaCl (0-15% w/v) and effect of pH (8-11) on the production of antimicrobial compound was checked. All the flasks were incubated at 30°C for 7-8 days. Then the cell free filtrates of the culture were collected from all the flasks separately and were tested against the actively growing sensitive culture of *S. flexneri* by agar well diffusion method.

### 2.5 Production and Extraction of Antimicrobial Metabolites

The strain *Nocardiopsis sp.GhM-HA-6* was cultivated in starch broth with 10% w/v NaCl and pH 9 on a rotary shaker (120 rpm) at 37°C for 8 days. The cell-free extract was obtained by filtration of broth culture using Whatman No. 1 filter paper. Equal volume of ethyl acetate was added to the culture filtrate for the extraction of the bioactive compounds. The mixture was added to 250 ml glass flask, sealed with cotton plug, followed by aluminum foil to reduce evaporation of the organic solvent and placed on shaker for 2 hours at 150 rpm. Post agitation the mixture was transferred to a separating funnel to generate different layers; the organic layer that contained the secondary metabolites and the aqueous layer. The crude extract was obtained by concentrating the solvent by evaporation and stored at 4°C for further use.

### 2.6 Identification of Antimicrobial Metabolites by Using Gas Chromatography-Mass Spectrometry (GC-MS) Analysis

Identification of the chemical compounds present in the crude ethyl acetate extract of *Nocardiopsis sp.GhM-HA-6* was carried out by Gas Chromatography-Mass Spectrometry (GC-MS) analysis. Analysis was conducted on a capillary column (Rxi-5ms, 30m, 0.25 mm id, 0.25 μm film thickness) with the following conditions: constant flow of Helium, 1.0 ml min^−^1; the fixed inlet temperature, 285°C throughout the analysis; injection volume, 3 μl in the linear with an open purge valve (30:1 split ratio); Linear velocity: 36.8 cm/second; Pressure: 65.0 kPa; Purge flow: 3.5 ml/min; Column flow: 1 ml/min; Oven ramp: 80°C holds for 2.0 min, 18°C/min to 260°C holds for 6.0 min, 4°C/min to 285°C holds for 6.0 min; Total run time: 30.25 min. The MS instrument with Ion source temperature: 200°C; Interface temperature: 300°C; Solvent cut time: 5.0 min; Detector voltage: 1 kV; Acquisition mode: Scan mode; Scan speed: 909; Event time: 0.78 second; starting m/z: 40 to 700 m/z. The peaks were identified by comparing the mass spectra with the National Institute of Standards and Technology (NIST, USA) library.

## 3. Results

### 3.1 Isolation and Morphology of the Organism

*Nocardiopsis sp. GhM-HA-6*, a haloalkaliphilic actinomycete was isolated from the Gulf of Khambhat, Western India. The isolate was characterized on the basis of its cell and colony morphology and Gram’s reaction. The colonies were small sized, round shaped, regular, slightly raised, rough, and opaque. It was Gram-positive, having a filamentous, long thread-like structure. It started sporulation on starch agar after 3 days of incubation with a fluffy mass of spores.

### 3.2 Molecular Identification of Nocardiopsis sp. GhM-HA-6 through 16S rRNA Sequencing

The 16S rRNA gene sequencing of the strain *Nocardiopsis sp. GhM-HA-6* showed the presence of 1456 bp long 16S rRNA gene in the genomic sequence. The sequence was submitted to NCBI, GenBank, Maryland, USA with accession number (KF384492). The phylogenetic tree was constructed using the neighbor-joining with Kimura 2-state parameter and pairwise-deletion model analysis implemented in the program MEGA software version X (Kumar et. al., 2018) and also evaluated in a bootstrap analysis of 1,000 replicates (Figure 1). The molecular characterization through 16S rRNA gene sequencing it was revealed that the strain belonged to *Nocardiopsis sp.*

### 3.3 The Antimicrobial Potential

The antimicrobial potential of the isolate was checked against six Gram-positive, eight Gram-negative bacteria and three fungi. The isolate *Nocardiopsis sp. GhM-HA-6* inhibited the growth of five Gram-positive bacteria *Bacillus subtilis, Staphylococcus aureus, Micrococcus luteus, Staphylococcus epidermidis, Bacillus cereus*, and three Gram-negative bacteria *Shigella flexneri, Pseudomonas aeruginosa, Serratia marcescens* (Figure 2). The highest inhibition was recorded against *Shigella flexneri*, however the isolate didn’t show any inhibitory effect against fungi.

### 3.4 Optimization of Growth Conditions for Antimicrobial Compound Production

To achieve the highest production of antimicrobial compounds, effect of various growth conditions were studied. Optimum production of the antimicrobial compound, against *S. flexneri,* was obtained in starch broth followed by soybean malt broth, starch casein broth and actinomycetes broth. However, *Nocardiopsis sp. GhM-HA-6* wasn’t able to produce any kind of antimicrobial compounds in complete broth, glucose asparagine broth and glucose glycine broth (Figure 3). Optimization of various growth conditions showed that the highest antimicrobial compound production was obtained in the presence of 0.5 % w/v starch, 1% w/v Yeast extract, 10% w/v NaCl and pH 9 (Figure 4-7).

### 3.5 Identification of Bioactive Compounds by GC-MS Analysis of Ethyl Acetate Extract

Identification of the bioactive compounds, present in ethyl acetate extract, was carried out using GC-MS analysis. The GC-MS chromatogram of the *Nocardiopsis sp. GhM-HA-6* crude extract showed a total of 18 peaks (Figure 8). When compared with the NIST database, the nearest compound hits for those peaks were found (Table 1). The compounds identified included Phenol, 2, 4-bis (1, 1-dimethylethyl)-, various alkanes (Heptadecane, Heneicosane, Docosane, Tricosane, Tetracosane, Pentacosane, Hexacosane, Heptacosane, Tetratetracontane, Octacosane, Nonacosane, Triacontane, Hentriacontane) and methyl derivatives of alkane (2-methylpentacosane, 2-methylhexacosane, 2, 24-dimethylpentacosane, 2-methylheptacosane).

**Table 1:** GC-MS analysis of *Nocardiopsis sp.* GhM-HA-6 ethyl acetate extract

| Peak No. | Retention Time | Name of the Compound | Molecular Formula | Molecular Weight | Peak Area % | Reference |
| --- | --- | --- | --- | --- | --- | --- |
| 1 | 8.478 | Phenol, 2,4-bis(1,1-dimethylethyl)- | C<sub>14</sub>H<sub>22</sub>O | 206.32 | 0.52 | Padmavathi AR et. al., 2014;<br>Nandhini US et. al., 2015 |
| 2 | 10.980 | Heptadecane | C<sub>17</sub>H<sub>36</sub> | 240.46 | 1.09 | Faridha Begum I et. al., 2016 |
| 3 | 11.892 | Heneicosane | C<sub>21</sub>H<sub>44</sub> | 296.6 | 1.76 | Nandhini US et. al., 2015 |
| 4 | 12.413 | Docosane | C<sub>22</sub>H<sub>46</sub> | 310 | 5.85 | El-Naggar NE et. al., 2017 |
| 5 | 12.990 | Tricosane | C<sub>23</sub>H<sub>48</sub> | 324 | 13.77 | El-Naggar NE et. al., 2017 |
| 6 | 13.656 | Tetracosane | C<sub>24</sub>H<sub>50</sub> | 338 | 20.94 | El-Naggar NE et. al., 2017 |
| 7 | 14.447 | Pentacosane | C<sub>25</sub>H<sub>52</sub> | 352 | 18.26 | El-Naggar NE et. al., 2017 |
| 8 | 15.026 | Hexacosane | C<sub>26</sub>H<sub>54</sub> | 366 | 1.35 | El-Naggar NE et. al., 2017 |
| 9 | 15.408 | 2-methylpentacosane | C<sub>26</sub>H<sub>54</sub> | 366.7 | 14.08 | -- |
| 10 | 16.124 | Heptacosane | C<sub>27</sub>H<sub>56</sub> | 380 | 0.95 | El-Naggar NE et. al., 2017 |
| 11 | 16.264 | Tetratetracontane | C<sub>44</sub>H<sub>90</sub> | 619.2 | 1.05 | Kumar N et. al., 2019 |
| 12 | 16.603 | 2-methylhexacosane | C<sub>27</sub>H<sub>56</sub> | 380.7 | 8.28 | Kumar N et. al., 2019 |
| 13 | 17.503 | 2,24-dimethylpentacosane | C<sub>27</sub>H<sub>56</sub> | 380.7 | 0.56 | -- |
| 14 | 18.104 | 2-methylheptacosane | C<sub>28</sub>H<sub>58</sub> | 394.8 | 4.78 | -- |
| 15 | 19.356 | Octacosane | C<sub>28</sub>H<sub>58</sub> | 394 | 0.69 | El-Naggar NE et. al., 2017 |
| 16 | 19.797 | Nonacosane | C<sub>29</sub>H<sub>60</sub> | 408 | 2.52 | El-Naggar NE et. al., 2017 |
| 17 | 21.498 | Triacontane | C<sub>30</sub>H<sub>62</sub> | 422 | 1.52 | El-Naggar NE et. al., 2017 |
| 18 | 23.178 | Hentriacontane | C<sub>31</sub>H<sub>64</sub> | 436.85 | 1.01 | Faridha Begum I et. al.,2016 |

## 4. Discussion

The declining trend in the discovery of new antimicrobial compounds and the enigmatic development of antibiotic resistance in bacteria has increased the need for scientists to search for novel antimicrobial compounds. Microbial life can be found in the most diverse conditions, including extremes of temperature, pressure, salinity, pH, nutrient concentrations, water availability and presence of high levels of radiations, harmful heavy metals and toxic compounds such as organic solvents and hydrocarbons (Thumar and Singh, 2008). Ability to sustain under extremity makes them interesting system to study the biochemical and molecular basis of adaptations, besides biotechnological aspects (Sharma *et. al*., 2012). In the light of this knowledge, we should explore new habitats in order to identify novel actinobacterial isolates and bioactive compounds. Recently, extremophilic actinobacteria have attracted the researchers with the hope that these organisms would add an inventive dimension to naturally available antimicrobial products research (Zitouni et. al., 2004; Vijayakumar et. al., 2012; Dhanasekaran et. al., 2014). Significant studies have not been conducted so far in the Gulf of Khambhat. Therefore, the present study was intended to isolate, screen and characterize antibiotics producing haloalkaliphilic actinomycetes from the Gulf of Khambhat. *Nocardiopsis sp. GhM-HA-6*, a haloalkaliphilic actinomycete was isolated from the saline soil sample collected from Ghogha site of Bhavnagar district. The isolate was identified as Gram-positive, filamentous, long thread-like structure containing, spore forming actinomycete based on cell and colony morphology and Gram’s staining while molecular characterization confirmed the isolate as *Nocardiopsis sp. GhM-HA-6.* The isolate showed the highest similarity with *Nocardiopsis salina* YIM 90010 (98.46%) and the lowest similarity with *Actinomadura gamaensis* (90.61%). The isolate was able to inhibit the growth of total eight test organisms. The antimicrobial compounds produced by the isolate were more effective against Gram-positive bacteria as compared to Gram-negative bacteria while the isolate didn’t show any inhibitory effect against fungi. Among all selected test organisms, the highest sensitivity was observed against *Shigella flexneri*, the intestinal tract pathogen. Our results are quite comparable with Sharma *et. al.*, (2016) who checked the broad-spectrum antimicrobial activity of forest derived soil Actinomycete, *Nocardia sp. PB-52*. Nithya *et. al*., (2018) isolated 134 morphologically distinct actinobacteria from various soil samples collected from the Saudi Arabian desert. They also used the same methods for isolation, identification, and characterization of antimicrobial compounds from the actinomycete isolates. Range of parameters can affect the production of antimicrobial compounds, so to increase the yield, effect of growth media, carbon source, nitrogen source, NaCl and pH was checked on the production of antimicrobial compounds by isolate. Optimization of media showed that the highest antimicrobial compound production and the highest antimicrobial activity against *S. flexneri* was found in starch broth followed by soybean malt broth, starch casein broth and actinomycetes broth. Optimization showed that the higher production of antimicrobial compounds was observed in media supplemented with complex carbon sources such as starch and malt while production of antimicrobial compounds wasn’t found in media supplemented with glucose like simple carbon source. So for further production processes starch was used as a carbon source. The optimization of starch concentration indicated that 0.5% w/v is the optimum concentration required for the highest production of antimicrobial compounds, further increase or decrease in concentration of starch inhibits the production of antimicrobial compounds. Yeast extract was provided as the nitrogen source, initial increase in concentration of yeast extract increased the production of antimicrobial compounds upto 1% w/v, further increase in yeast extract concentration decreased the production. The isolate showed the highest production of antimicrobial compounds at 10% w/v NaCl followed by 15 % w/v NaCl. The isolate wasn’t able to produce antimicrobial compounds in absence or lower amount of NaCl. Optimization of pH showed that the isolate was able to produce antimicrobial compounds at pH 9 and 10 but the highest activity was observed at pH 9. Identification of the bioactive compounds, present in ethyl acetate extract, was carried out using GC-MS analysis. The GC-MS chromatogram of the *Nocardiopsis sp. GhM-HA-6* ethyl acetate extract indicated the presence of 18 compounds, from which 15 compounds including Phenol, 2, 4-bis (1, 1-dimethylethyl)- and various types of alkanes have already been reported to have antimicrobial potential against variety of pathogens (Padmavathi et. al., 2014; Nandhini et. al., 2015; Faridha Begum et. al., 2016; El-Naggar et. al., 2017; Kumar et. al., 2019) while methyl derivatives of alkanes identified from ethyl acetate extract haven’t been reported for antimicrobial activity. Padmavathi et. al., (2014) reported Phenol-2, 4-bis (1, 1-dimethylethyl) – derived from marine bacteria as an efficient antimicrobial agent against uropathogen *Serratia marcescens.* Recently, the study conducted by Kumar *et. al.*, (2014c), showed the highest antimicrobial activity in the GC-MS fractions containing the highest amount of phenolic compounds. In addition, 3, 5-bis (1, 1-dimethylethyl)-phenol and 2, 4-di-t-butyl-6-nitrophenol were the two phenolic compounds detected in EA-PB-52. Roy *et. al.*, (2015) also reported a surfactant activity of 3, 5-bis (1, 1-dimethyethyl)-phenol in *Nocardiopsis* VITSISB isolated from Marina beach, India.

## 4. Conclusion

Literature studies say that the discovery of new antibiotics remains one of the most important tasks of modern biotechnology due to the rapid emergence of antibiotic resistance among pathogenic bacteria. To meet the search for novel antimicrobial compounds, the present study was intended to explore untapped coastal region of Gulf of Khambhat. Ghogha coast was selected because of its unique geographical location with respect to the diversity, bioactive potential of actinomycetes. *Nocardiopsis sp.* GhM-HA-6, isolated from the same site, inhibited the growth of variety of test organisms including *Staphylococcus epidermidis, Pseudomonas aeruginosa, Shigella flexneri,* and *Staphylococcus aureus*. The ethyl acetate extract of *Nocardiopsis sp.* GhM-HA-6, indicated the presence of potent antimicrobial compounds such as Phenol-2, 4-bis (1, 1-dimethylethyl)- which can further be purified and used for the treatment of various known and unknown infections. In nutshell, our findings in this field will surely be a significant contribution to the knowledge of compounds unique from marine bacteria as potential sources of new drugs in the pharmacological industry.

## 5. Acknowledgement

Financial Assistance from the Department of Biotechnology (New Delhi, India) is acknowledged.

---

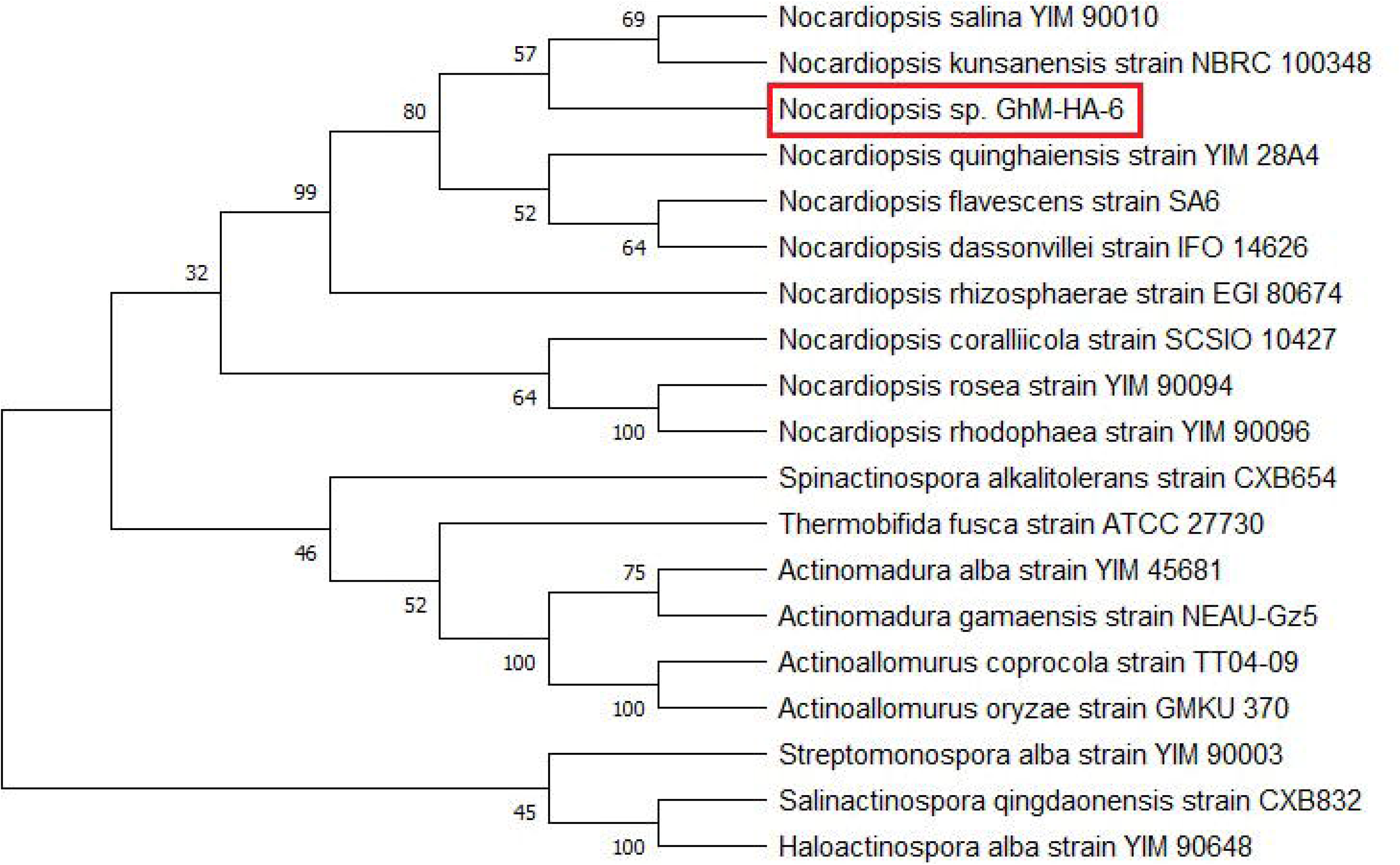

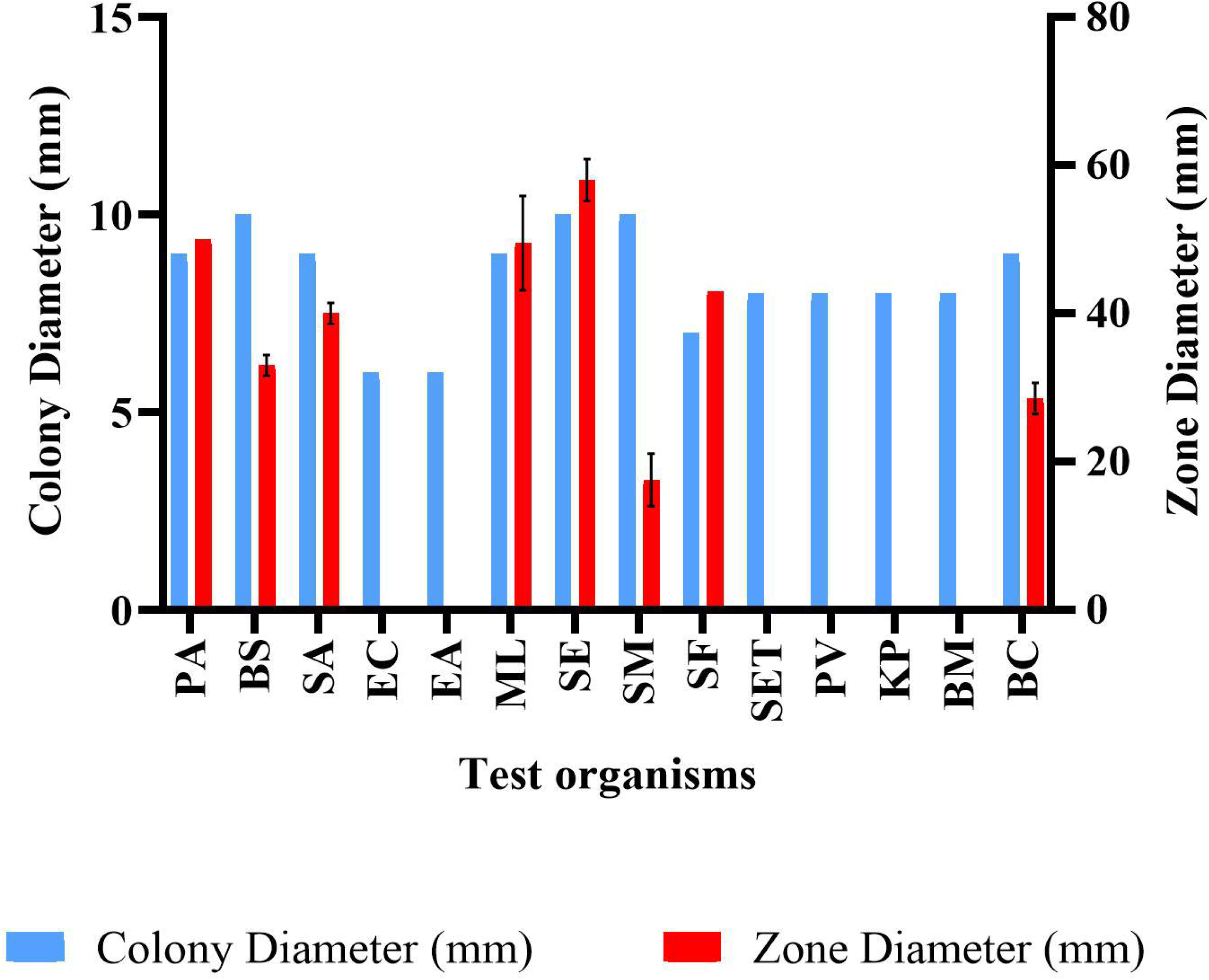

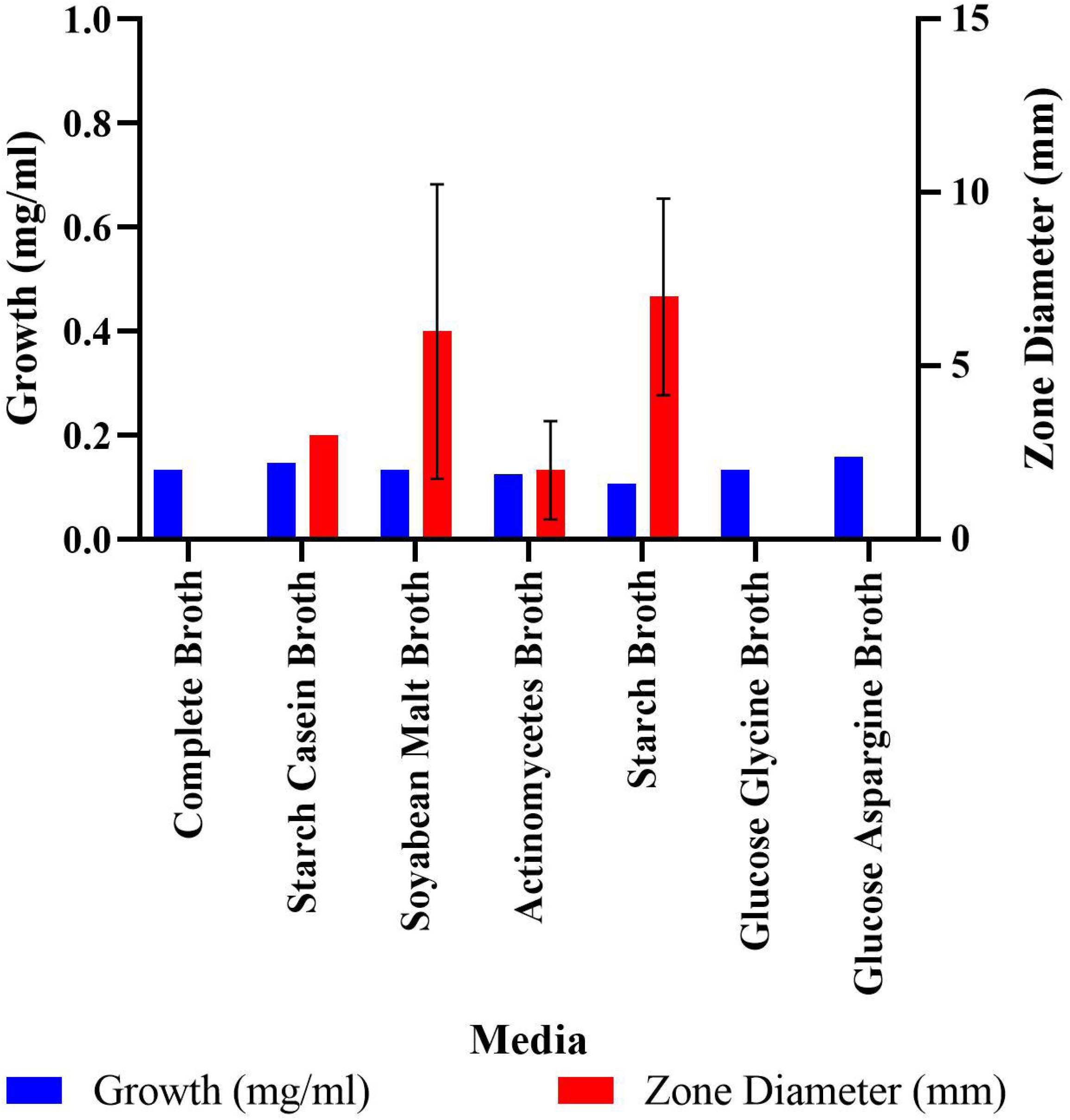

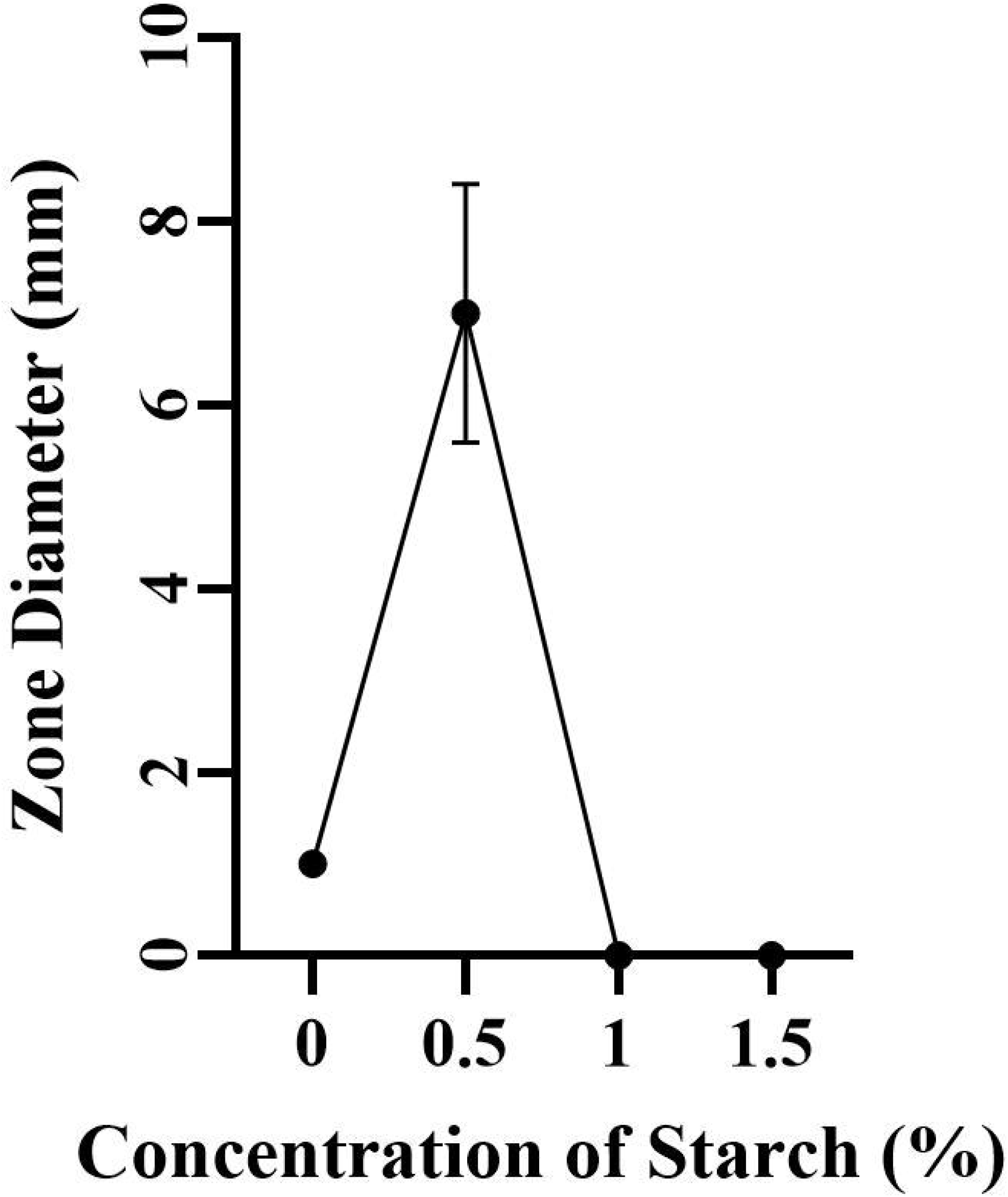

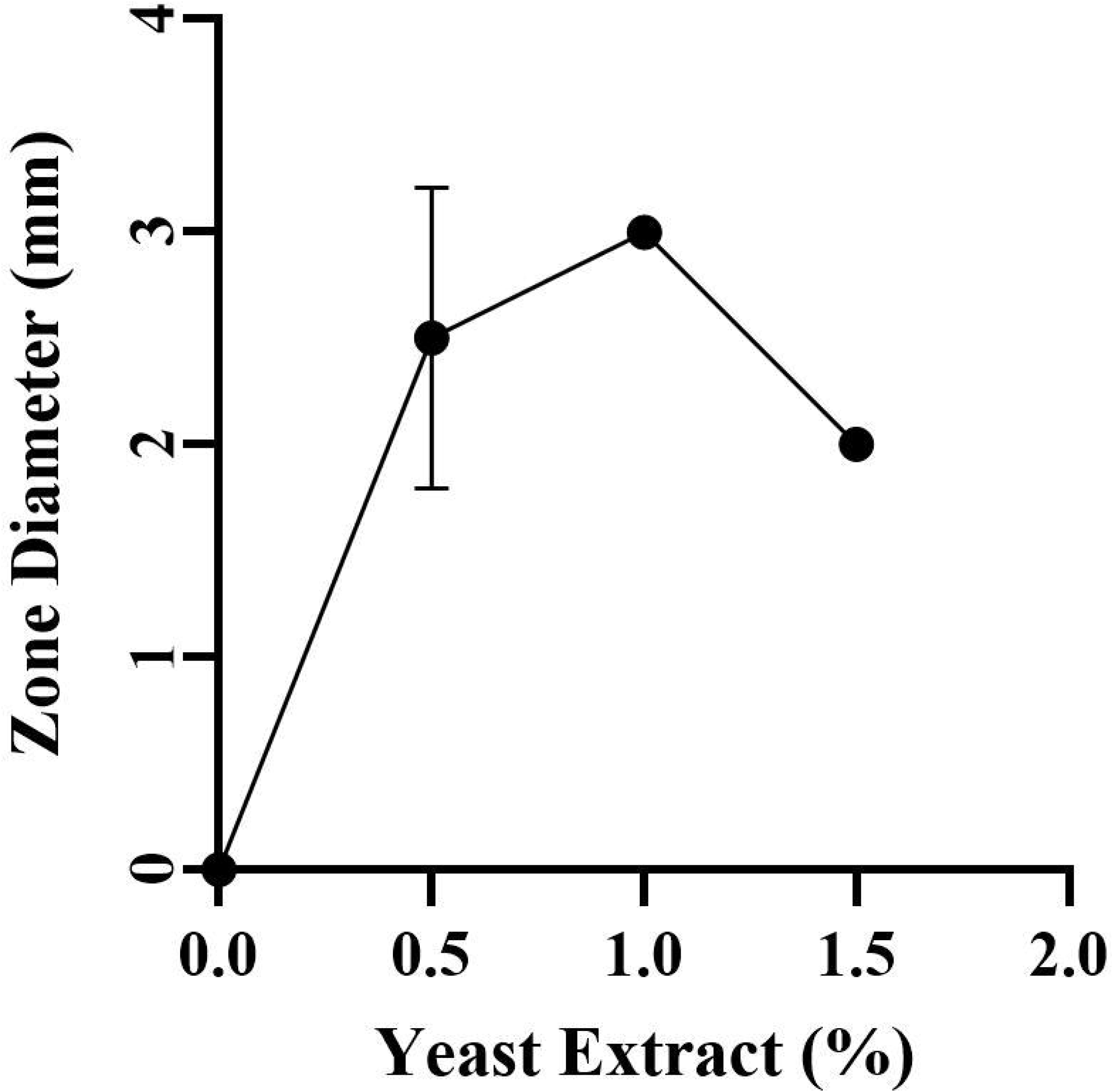

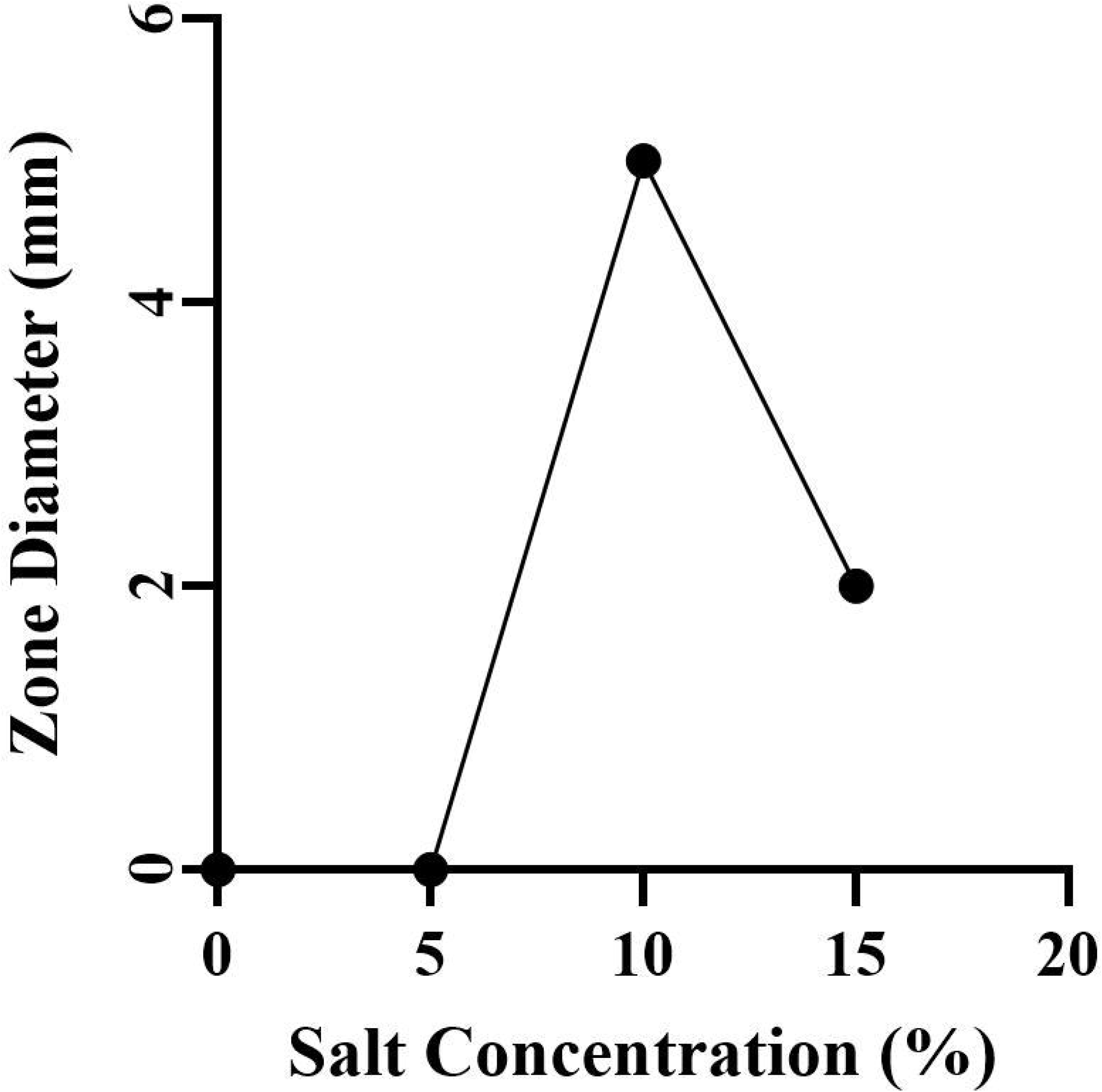

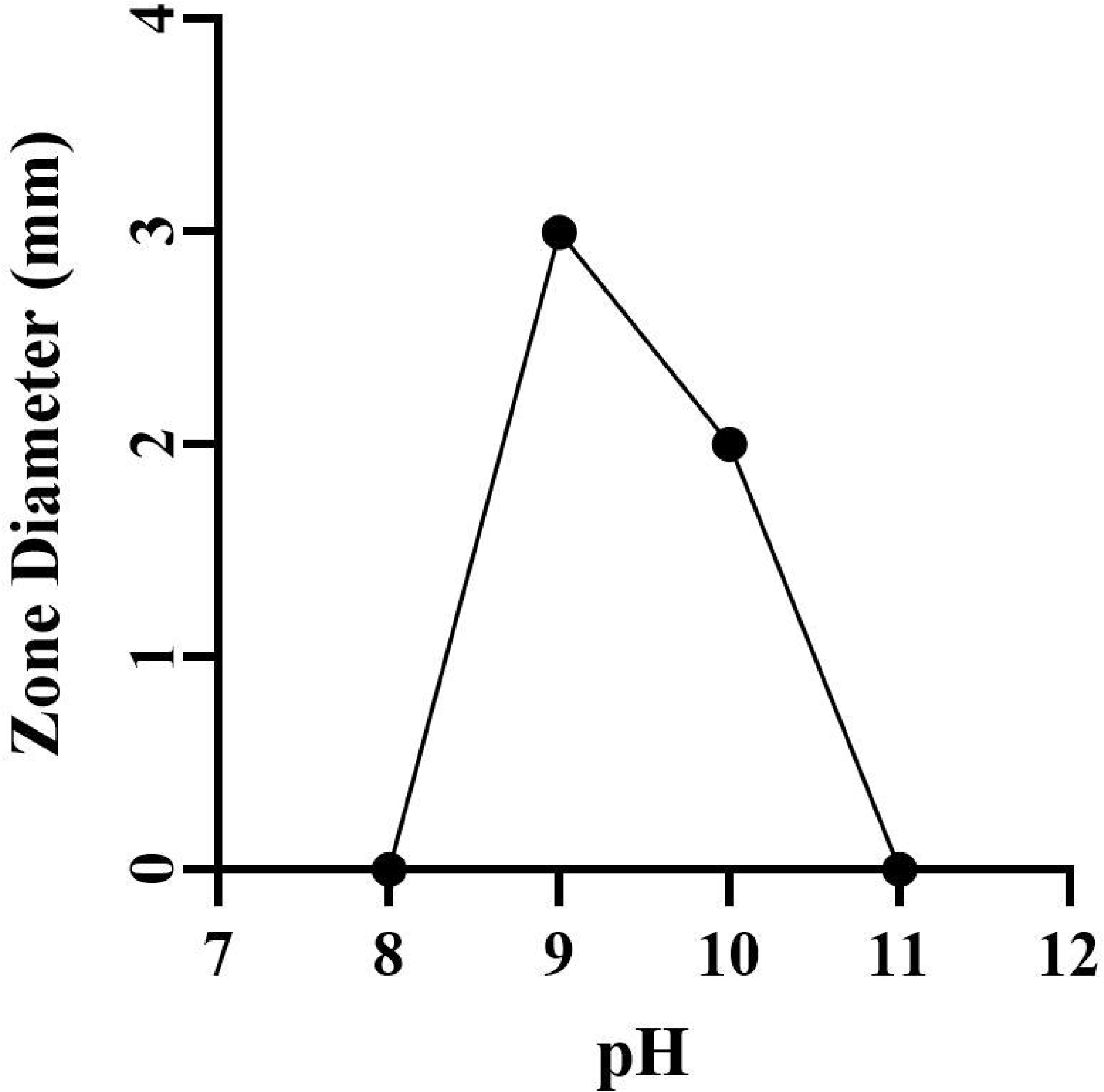

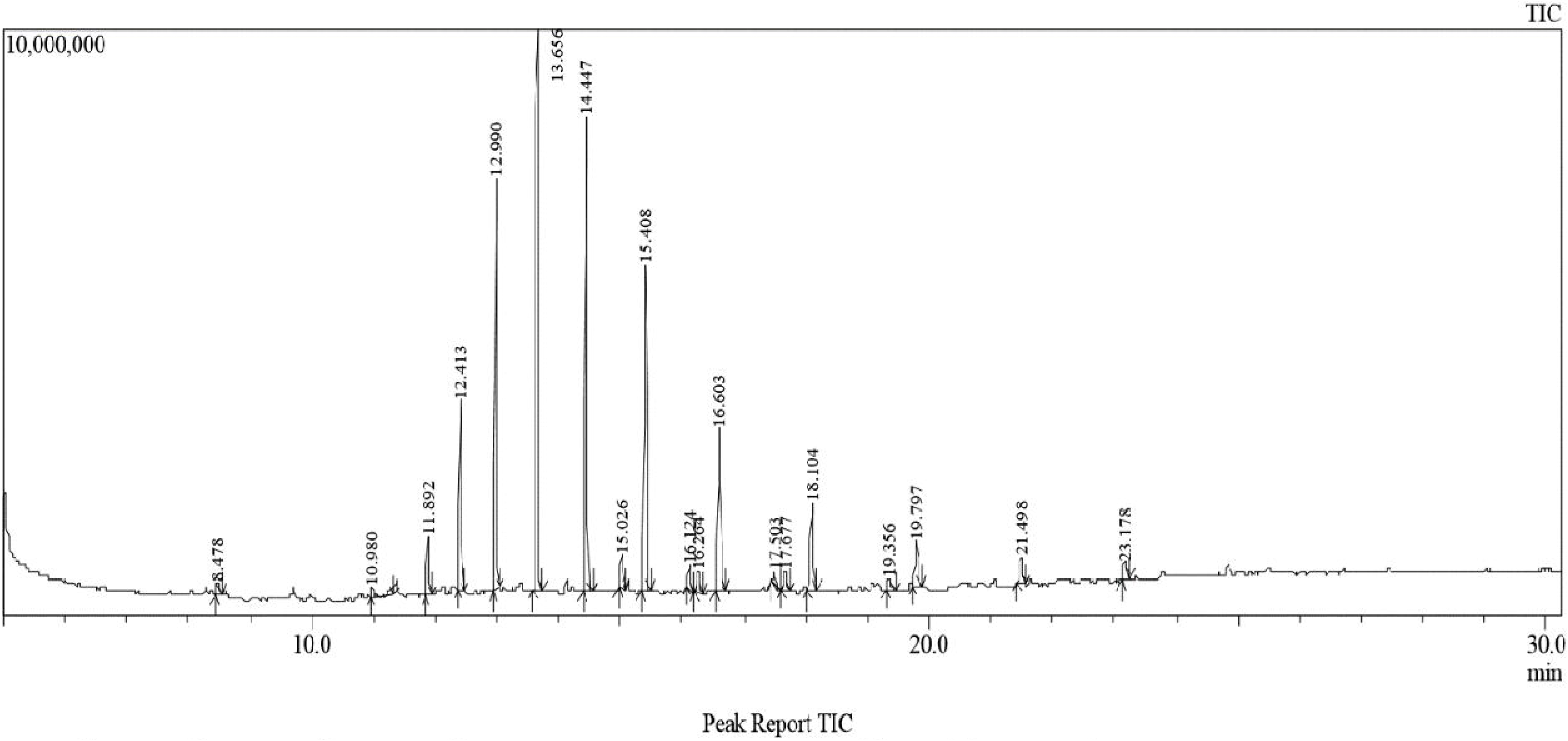

## References

1. Becerril-Espinosa A, Freel KC, Jensen, PR and Soria-Mercado IE, 2013. Marine Actinobacteria from the Gulf of California: diversity, abundance and secondary metabolite biosynthetic potential. Antonie van Leeuwenhoek 103, 809–819. https://doi.org/10.1007/s10482-012-9863-3

2. Bennur T, Kumar AR, Zinjarde S and Javdekar V, 2015. *Nocardiopsis* species: Incidence, ecological roles and adaptations. Microbiological Research, 174, 33–47 https://doi.org/10.1016/j.micres.2015.03.010

3. Chakrabarti T, 1998. ACTINOMYCETES – Isolation, Screening, Identification, and Gene Cloning in Streptomyces Laboratory manual.

4. Cook AE and Meyers PR, 2003. Rapid identification of filamentous actinomycetes to the genus level using genus-specific 16S rRNA gene restriction fragment patterns. Int J Syst Evol Microbiol, Nov; 53 (Pt 6): 1907–15. doi: https://doi.org/10.1099/ijs.0.02680-0 PMID: 14657122.

5. Dhanasekaran D, Thajuddin N, Panneerselvam A and Chandraleka S, 2014. Isolation, Characterization of Antibacterial Methyl Substituted β-Lactum Compound from *Streptomyces noursei* DPTD21 in Saltpan Soil, India. TBAP, 4 (2) 71 – 88.

6. Dolak LA, Castle TM and Laborde AL, 1980. 3-Trehalosamine, a new disaccharide antibiotic. J Antibiot, 7:690–4.

7. El-Naggar NE, El-Bindary AA, Abdel-Mogib M and Nour NS, 2017. In vitro activity, extraction, separation and structure elucidation of antibiotic produced by *Streptomyces anulatus* NEAE-94 active against multi-drug resistant *Staphylococcus aureus*, Biotechnology & Biotechnological Equipment, 31:2, 418–430, DOI: https://doi.org/10.1080/13102818.2016.1276412

8. Engelhardt K, Degnes KF, Kemmler M, Bredholt H, Fjaervik E, Klinkenberg G, Sletta H., Ellingsen TE and Zotchev SB, 2010a. Production of a new thiopeptide antibiotic TP-1161, by a marine *Nocardiopsis species*. Appl Environ Microbiol; 76:4969–76.

9. Faridha Begum I, Mohankumar R, Jeevan M and Ramani K, 2016. GC-MS Analysis of Bio-active Molecules Derived from *Paracoccus pantotrophus* FMR19 and the Antimicrobial Activity Against Bacterial Pathogens and MDROs. Indian J Microbiol. 2016 Dec; 56(4):426–432. doi: https://doi.org/10.1007/s12088-016-0609-1 Epub, Jul 20. PMID: 27784938; PMCID: PMC5061700.

10. Jose PA and Jebakumar SR, 2012. Phylogenetic diversity of actinomycetes cultured from coastal multipond solar saltern in Tuticorin, India. Aquat Biosyst, 8:23.

11. Kroppenstedt RM and Evtushenko LI, 2006. The family *Nocardiopsaceae*. The Prokaryotes: A Handbook on the Biology of Bacteria, In: Dworkin M, Falkow S, Rosenberg E, Schleifer KH, Stackebrandt E, editors., New York, New York: Springer-Verlag, vol. 3, 3rd ed.; p. 754–95.

12. Kumar N, Menghani E and Mithal R, 2019. GCMS analysis of compounds extracted from Actinomycetes AIA6 isolates and study of its antimicrobial efficacy, Indian Journal of Chemical Technology, Vol. 26, July 2019, 362–370.

13. Kumar N, Singh RK, Mishra SK, Singh AK and Pachouri UC, 2010. Isolation and screening of soil Actinomycetes as source of antibiotics active against bacteria. International Journal of Microbiology Research, Volume 2, Issue 2, 2010, pp-12–16, ISSN: 0975-5276.

14. Kumar PS, Duraipandiyan V and Ignacimuthu S, 2014. Isolation, screening and partial purification of antimicrobial antibiotics from soil *Streptomyces sp.* SCA 7. Kaohsiung Journal of Medical Sciences, 30 (9): 435–446.

15. Kumar S, Stecher G, Li M, Knyaz C and Tamura K. (2018). MEGA X: Molecular Evolutionary Genetics Analysis across Computing Platforms. Mol. Biol. Evol., 35, 1547–1549.

16. Meklat A, Sabaou N, Zitouni A, Mathieu F, Lebrihi A, 2011. Isolation, taxonomy, and antagonistic properties of halophilic actinomycetes in Saharan soils of Algeria. Appl Environ Microbiol; 77:6710–4.

17. Meyer J, 1976. *Nocardiopsis,* a new genus of the order Actinomycetales. Int J Syst Bacteriol, 26:487–93.

18. Mwirichia R, Muigai AW, Tindall B, Boga HI and Stackebrandt E, 2010. Isolation and characterization of bacteria from the haloalkaline Lake Elmenteita, Kenya. Extremophiles, 14:339–48.

19. Nandhini US, Sangareshwari S and Lata K, 2015. Gas chromatography-mass spectrometry analysis of bioactive constituents from the marine *Streptomyces*. Asian J Pharm Clin Res, 8:244–246.

20. Nithya K, Muthukumar C, Biswas B, Alharbi NS, Kadaikunnan S, Khaled JM and Dhanasekaran D, 2018. Desert actinobacteria as a source of bioactive compounds production with a special emphasis on Pyridine-2, 5-diacetamide a new pyridine alkaloid produced by *Streptomyces sp. DA3-7*. Microbiological Research, 207, 116–133.

21. Padmavathi AR, Bose A and Pandian SK, 2014. Phenol, 2, 4-bis (1, 1-dimethylethyl) of marine bacterial origin inhibits quorum sensing mediated biofilm formation in the uropathogen *Serratia marcescens*. Biofouling, Vol. 30, No. 9, 1111–1122, https://doi.org/10.1080/08927014.2014.972386

22. Rainey FA, Rainey NW, Kroppenstedt RM and Stackebrandt E, 1996. The Genus *Nocardiopsis* represents a phylogenetically coherent taxon and a distinct Actinomycete lineage: proposal of *Nocardiomaceae* fam. nov. Int J Syst Bacteriol, 46:1088–92.

23. Ramnathan V, Vincent D, Sundrmoorthy S, Shunmungraj T, 2002. Critical habitat information system for Gulf of Khambhat-Gujarat; Govt. of India: Department of Ocean development.

24. Romanenko LA, Tanaka N, Kalinovskaya NI and Mikhailov VV, 2013. Antimicrobial potential of deep surface sediment associated bacteria from the Sea of Japan. World J Microbiol Biotechnol; 29: 1169–77.

25. Roy S, Chandni S, Das I, Karthik L, Kumar G, and Rao KVB, 2015. Aquatic model for engine oil degradation by rhamnolipid producing *Nocardiopsis* VITSISB. Biotech, 5, 153–164. doi: https://doi.org/10.1007/s13205-014-0199-8

26. Sharma AK, Gohel S and Singh SP, 2015. Actinobase: Database on molecular diversity, phylogeny and biocatalytic potential of salt tolerant alkaliphilic actinomycetes. Bioinformation; 8 (11): 535–538. DOI: https://doi.org/10.6026/97320630008535

27. Sharma P, Kalita MC, and Thakur D, 2015. Broad Spectrum Antimicrobial Activity of Forest-Derived Soil Actinomycete, *Nocardia sp. PB-52*. Frontiers in microbiology, 7, 347. https://doi.org/10.3389/fmicb.2016.00347

28. Sivakumar K, Sahu MK, Thangaradjou T and Kannan L, 2007. Research on marine actinobacteria in India. Indian J. Microbiol., 47: 186–196 https://doi.org/10.1007/s12088-007-0039-1

29. Solanki D, Kanejiya J, Beleem I and Gohil B, 2016. Checklist of intertidal marine fauna in mangrove ecosystem, Ghogha coast, Gulf of Khambhat, India. Journal of Entomology and Zoology Studies; 4(4): 1281–1284.

30. Takahashi A, Hotta K, Saito N, Morioka M, Okami Y and Umezawa H, 1986. Production of novel antibiotic, dopsisamine, by a new subspecies of *Nocardiopsis mutabilis* with multiple antibiotic resistance. J Antibiotics; 39:175–83.

31. Tamura K, Dudley J, Nei M and Kumar S, 2007. MEGA 4: Molecular Evolutionary Genetics Analysis (MEGA) software version 4.0. Molecular Biology and Evolution, 24:1596–1599.

32. Thumar JT and Singh SP, 2008. Organic solvent tolerance of an alkaline protease from salt-tolerant alkaliphilic *Streptomyces clavuligerus* strain Mit-1, J Ind Microbiol Biotechnol, 6:211–218. DOI https://doi.org/10.1007/s10295-008-0487-6

33. Vijayakumar R, Panneer selvam K, Muthukumar C, Thajuddin N, Panneerselvam A and Saravanamuthu R, 2012. Antimicrobial potentiality of a halophilic strain of *Streptomyces sp. VPTSA18* isolated from the saltpan environment of Vedaranyam, India. Ann. Microbiol., 62, 1039–1047.

34. Wu ZC, Li S, Nam SJ, Liu Z and Zhang C, 2013. Nocardiamides A and B, two cyclohexapeptides from the marine-derived actinomycete *Nocardiopsis sp.* CNX037. J Nat Prod; 76:694–701.

35. Zitouni A, Boudjella H, Lamari L, Badji B, Mathieu F and Lebrihi A, 2005. *Nocardiopsis* and *Saccharothrix* genera in Saharan soils in Algeria: isolation, biological activities and partial characterization of antibiotics. Res Microbiol, 156:984–93.

36. Zitouni A, Lamari L, Boudjella H, Badji B, Sabaou N, Gaouar A, Mathieu F, Lebrihi A and Labeda DP, 2004. *Saccharothrix algeriensis sp. nov*.: isolated from Saharan soil. Int. J. Sys. Evol. Microbiol, 54, 1377–1381.

